# Fertility costs of cryptic viral infections in a model social insect

**DOI:** 10.1101/2021.05.04.442681

**Authors:** Abigail Chapman, Esmaeil Amiri, Bin Han, Erin McDermott, Olav Rueppell, David R Tarpy, Leonard J Foster, Alison McAfee

**Affiliations:** Department of Biochemistry and Molecular Biology, Michael Smith Laboratories, University of British Columbia, Vancouver, British Columbia, Canada; Department of Biology, University of North Carolina at Greensboro, Greensboro, North Carolina, USA; Department of Entomology and Plant Pathology, North Carolina State University, Raleigh, North Carolina, USA; Institute of Apicultural Research, Chinese Academy of Agricultural Sciences, Beijing, China; Department of Biological Sciences, University of Alberta, Edmonton, Alberta, Canada

## Abstract

Declining insect populations emphasize the importance of understanding the drivers underlying reductions in insect fitness. Here, we investigated viruses as a threat to social insect reproduction, using honey bees as a model species. We report that in a sample of N = 93 honey bee (*Apis mellifera*) queens from nine beekeeping operations across a wide geographic range, high levels of natural viral infection are associated with decreased ovary mass. We confirmed this finding in an independent sample of N = 54 queens. Failed (poor quality) queens displayed higher levels of viral infection, reduced sperm viability, smaller ovaries, and altered ovary protein composition compared to healthy queens. We experimentally infected queens with Israeli acute paralysis virus (IAPV) and found that the ovary masses of IAPV-injected queens were significantly smaller than control queens, demonstrating a causal relationship between viral infection and ovary size. Queens injected with IAPV also had significantly lower expression of vitellogenin, the main source of nutrition deposited into developing oocytes, and higher levels of heat-shock proteins (HSPs), which are part of the honey bee’s antiviral response. This work together shows that viral infections occurring naturally in the field are compromising queen reproductive success.

## Introduction

Amidst a backdrop of widespread insect declines^1-6^ and fluctuating populations^7,8^, it is vitally important to better understand the impacts of interacting biotic and abiotic stressors on insect physiology^9^. Pesticide exposure and climate change are often cited as drivers of insect decline^9-13^, but biotic drivers, such as viruses, are comparatively understudied, despite some viruses exhibiting broad host ranges with the potential for widespread impacts across species. Stressors that reduce fertility are particularly worrisome, especially for social insects, as damage to a single individual (the queen) can weaken the fitness of the entire colony. Additionally, extreme temperatures can have immunomodulatory effects in some insects^14,15^; therefore, the stressors of climate change and viral infections may actually interact to have synergistic effects, further underscoring the need to better understand the effects of both stressors^16^.

Viruses that affect managed species like honey bees (*Apis mellifera*) can infect a broad range of insect host species, creating the potential for pathogen spillover to native species pollinating in the same regions^17-24^. Infection with Israeli acute paralysis virus (IAPV) and Kashmir bee virus (KBV), which infect both *Apis* and non-*Apis* pollinators, results in slower colony start-up and a reduction in egg laying in *Bombus terrestris*^25^. Furthermore, high titres of deformed-wing virus (DWV) and *Varroa destructor* virus-1 (VDV-1, also known as DWV-B) are associated with extreme cases of ovarian degeneration in honey bee queens^26^, but there is a need for more research in this area.

Beyond the symptomatic impacts on insect health, virus infections also have the potential to indirectly reduce fecundity, even in the absence of overt symptoms, through a trade-off between reproductive ability and immune activation^27^. Reproduction and immune processes, whether induced by a pathogen or constitutively maintained, are both energetically demanding. Given a finite resource supply, an individual may not be able to sustain both fully at the same time^27^. Across multiple insect orders, mated female insects tend to have reduced immunity compared to their unmated counterparts^27^. Correspondingly, immune-challenged individuals frequently show a reduction in reproductive output measured as reduced overall fecundity^28^, reduced protein in both ovaries and eggs^29,30^, reduced oviposition rate^30,31^, and reduced viability of stored sperm^32,33^. This potential compromise is particularly relevant for a honey bee queen, whose physiology is tailored for laying eggs and little else; their ovaries make up about one third of their body mass, and they can lay >1000 eggs a day, which is roughly equivalent to their own body weight^34,35^. Additionally, a queen’s resource investment in ovary size is highly plastic and responds dramatically to external stressors like nutrient availability^35^. Thus, viral infections could negatively impact fertility either through direct effects of infecting reproductive tissue or compromised allocation of resources associated with systemic infections. We have previously shown that lysozyme, an immune effector, is negatively correlated with stored sperm viability in honey bee queens and that failing queens, even without symptoms of viral infection, have significantly lower sperm viability and higher titers of black queen cell virus (BQCV) and sacbrood virus (SBV)^36^. This suggests that reproduction and immune activation, as a result of pathogenic challenge, are negatively associated in honey bee queens, at least in terms of sperm maintenance.

Here, we investigated the relationship between viral infection and reproductive physiology in honey bees, a model social insect. We measured the ovary masses of queens rated as ‘failed’ and ‘healthy’ by beekeepers and which had natural variations in viral infection. In two independent populations, we identified and validated a significant negative relationship between viral infection and ovary size. We measured ovarian protein investment associated with viral infection using quantitative proteomics, confirmed the upregulation of antiviral proteins in virus infected queens, and addressed the possibility of abiotic stressors as alternate explanations for our results. Finally, we have confirmed the causal, rather than correlative, effect of virus infection on ovaries by experimentally infecting queens with IAPV. This work suggests that pathogenic infections negatively impact reproductive quality and fitness in a model insect, supporting the idea that infection may be a significant driver of declining insect health.

## Results

### Failed queens have smaller ovaries and reduced sperm viability

In order to investigate if small ovary size is a general feature of queens with poor reproductive output (described by beekeepers as “failing”), we compared the ovary masses of failing and healthy queens collected from beekeeping operations in three independent queen surveys (denoted as Field Surveys 1-3) conducted in multiple geographical locations (described in **Table 1**). We found a highly significant reduction in ovary mass (linear mixed model, family = Gaussian; t = -4.37, df = 104, p < 0.0001) in failed queens across surveys (**Supplementary Table S1 and Figure 1a**), indicating that small ovaries are a widespread phenotype associated with poor fecundity, as described by beekeepers.

**Table 1.**
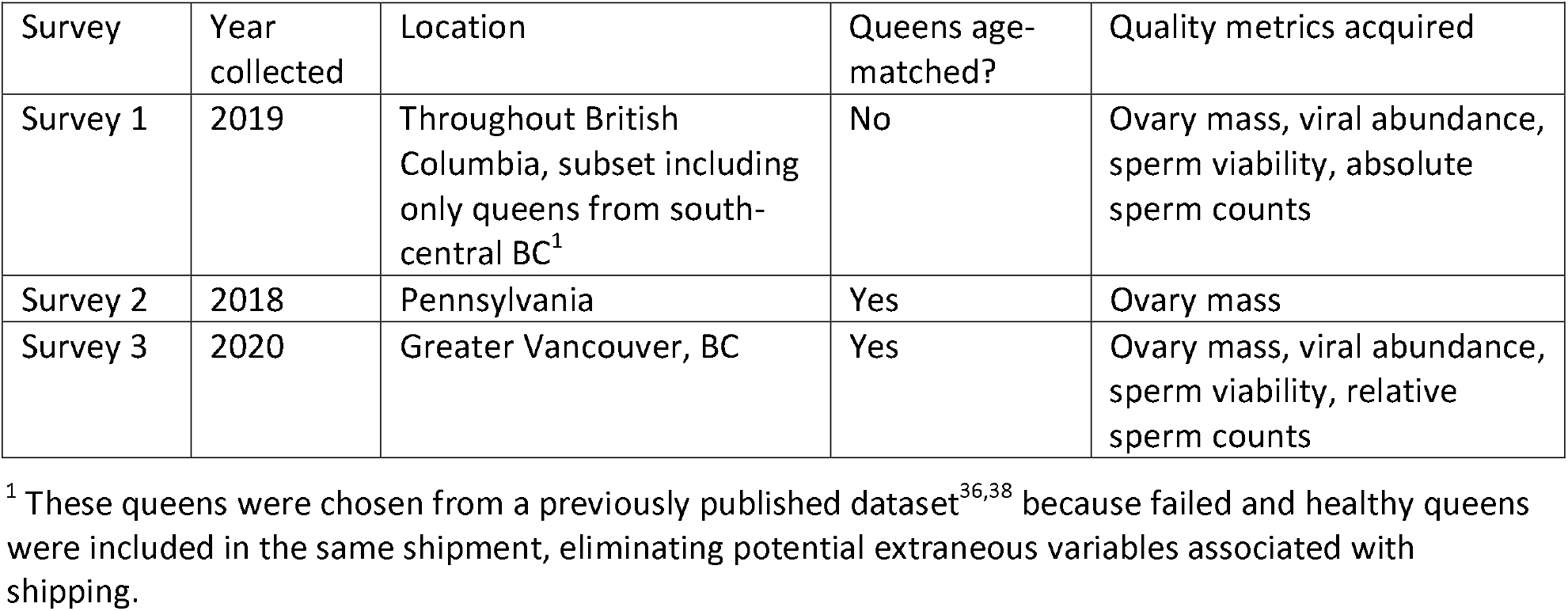
Summary of queen surveys

| Survey | Year collected | Location | Queens age-matched? | Quality metrics acquired |
| --- | --- | --- | --- | --- |
| Survey 1 | 2019 | Throughout British Columbia, subset including only queens from south-central BC<sup>1</sup> | No | Ovary mass, viral abundance, sperm viability, absolute sperm counts |
| Survey 2 | 2018 | Pennsylvania | Yes | Ovary mass |
| Survey 3 | 2020 | Greater Vancouver, BC | Yes | Ovary mass, viral abundance, sperm viability, relative sperm counts |

<sup>1</sup> These queens were chosen from a previously published dataset<sup>36,38</sup> because failed and healthy queens were included in the same shipment, eliminating potential extraneous variables associated with shipping.

**Figure 1.**
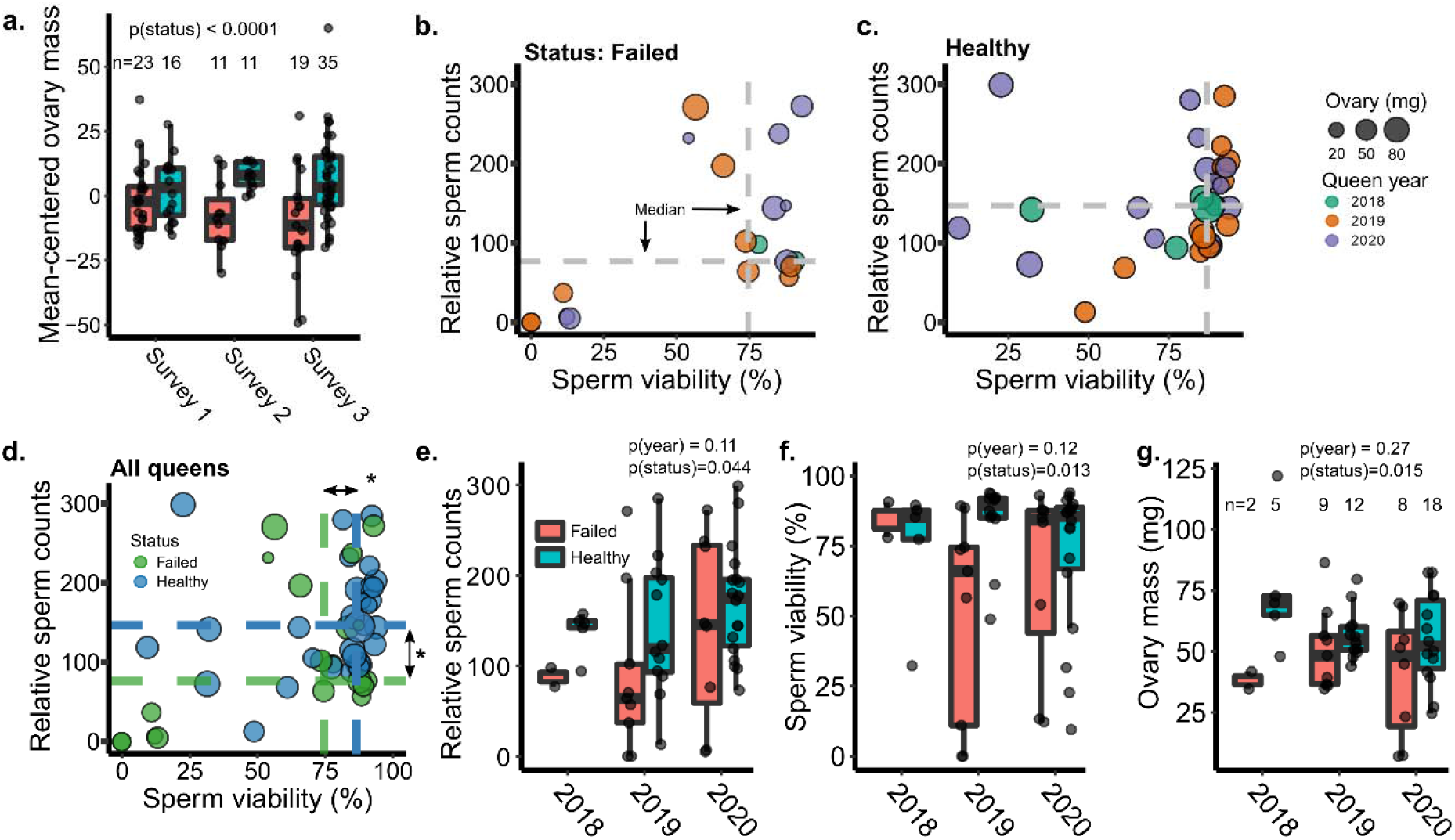
Fertility metrics of failed and healthy queens. In all cases, years indicate the year the queen was reared, and all queens were collected in the summer of 2020. a) Ovary masses of failed and healthy queens collected across three different surveys conducted in British Columbia and Pennsylvania (analyzed a linear mixed model with status as a fixed effect and source location as a random effect). Because the average ovary size differed between surveys, data were mean-centered by survey prior to analysis to better highlight the effect of status. Boxes represent the interquartile range, bars indicate the median, and whiskers span 1.5 times the interquartile range. b) Queens rated as ‘failed’ (spotty brood pattern, drone layer, or dwindling adult population) or c) ‘healthy’ (contiguous worker brood patterns, medium-strong adult population) by local beekeepers were collected in the summer of 2020. Sperm viability and sperm counts were determined by fluorescent imaging, and wet ovary weight was measured on an analytical balance. d) Statistical analyses on data presented in b) and c) were conducted using either a linear mixed model (ovary mass and sperm counts) or a generalized linear mixed model fitted by maximum likelihood (sperm viability; see **Table 2** for details). In the statistical models, queen age (0, 1, or 2 years, which corresponds to queens reared in 2020, 2019, and 2018, respectively) and health status (healthy or failed) were included as fixed effects and source location was included as a random effect. Asterisks indicate statistical significance (p < 0.05), with exact p values given in panels e-g. e-g) Same data as in d) but separated by the year in which the queen was reared (*i.e*., a 2018 queen was 2 years old).

In our previous queen survey, we also found that queen failure was associated with reduced stored sperm viability and sperm counts (not recorded for Survey 2)^36^. However, the ages of failed queens were unknown, leading to questions over whether differences in queen quality metrics were linked to queen failure or old age. We therefore collected an additional n = 54 queens (19 failed and 35 healthy) with known ages, the oldest of which were 2 years old, all from our local region to avoid extraneous effects of shipping. We found that sperm viability (generalized linear mixed model, family = binomial; z = -2.49, df = 45, p = 0.013) and relative sperm counts (linear mixed model, family = Gaussian; t = -2.08, df = 45, p = 0.044) were again significantly lower in failed queens, even when accounting for age as a fixed factor and apiary location as a random effect (**Figure 1b-f**), corroborating our previous studies^36,37^. See **Table 2** for complete statistical reporting and **Supplementary Table S2** for the underlying data. In this survey, failed queens also had significantly smaller ovaries (linear mixed model, t = -2.52, df = 45, p < 0.015) (**Figure 1g**).

**Table 2.** Statistical parameters

| Table 2. Statistical parameters |  |  |  |  |  |  |
| --- | --- | --- | --- | --- | --- | --- |
| Data set | Response | Model | Fixed effect | df | \|t/z/F\| | P |
| Ovary masses of queens in Field Surveys 1-3<sup>1</sup> | Ovary mass | Linear mixed model (lme) | Health status | 104 | 4.37 | < 0.0001 *** |
| Field Survey 1 queens (n = 93)<sup>2</sup> | Ovary mass | Linear mixed model (lme) | Total viral load (med-high) | 71 | 1.35 | 0.18 |
|  |  |  | Total viral load (low-high) | 71 | 2.24 | 0.028 * |
|  |  |  | Health status | 71 | 2.58 | 0.012 * |
| Field Survey 3 queens (n = 54)<sup>1</sup> | Ovary mass | Linear mixed model (lme) | Age | 43 | 0.24 | 0.81 |
|  |  |  | Health status | 43 | 2.16 | 0.036 * |
|  |  |  | Total viral load (low-high) | 43 | 1.78 | 0.082 |
|  |  |  | Total viral load (med-high) | 43 | 2.32 | 0.025 * |
|  | Sperm viability | Generalized linear mixed model (glmer) | Age | 43 | 0.48 | 0.63 |
|  |  |  | Health status | 43 | 1.78 | 0.075 |
|  |  |  | Total viral load (low-high) | 43 | 2.27 | 0.023 * |
|  |  |  | Total viral load (med-high) | 43 | 0.023 | 0.98 |
|  | Sperm counts | Linear mixed model (lme) | Age | 43 | 0.92 | 0.36 |
|  |  |  | Health status | 43 | 2.36 | 0.023 * |
|  |  |  | Total viral load (low- high) | 43 | 0.37 | 0.71 |
|  |  |  | Total viral load (med-high) | 43 | 0.91 | 0.37 |
| 2-week experimental queen stress | Ovary mass | Linear model (lm) | Pesticide | 23 | 1.49 | 0.25 |
|  |  |  | Temperature | 17 | 1.12 | 0.31 |
|  | Queen weight ratio | Linear model (lm) | Pesticide | 23 | 1.62 | 0.22 |
|  |  |  | Temperature | 17 | 1.65 | 0.12 |
| 2-day experimental queen stress | Ovary mass | Linear model (lm) | Treatment | 44 | 0.234 | 0.82 |

<sup>1</sup> Models included queen location as a random effect. Survey 2 contributed only ovary mass data, and not sperm metrics. Survey 2 data were analyzed in conjunction with ovary mass data from Surveys 1 and 3.
<sup>2</sup> Model did not include queen source location as a random effect due to singularities.

### Ovary size is inversely linked to viral abundance in naturally infected queens

Queen honey bees are susceptible to pathogenic infections^26,39-41^, and most commonly infected with sacbrood virus (SBV), black queen cell virus (BQCV), and deformed-wing virus (DWV)^36,39^. We previously found that the failed queens from Survey 1 have higher copy numbers of SBV and BQCV RNA^36^.We therefore hypothesized that ovary size is linked to viral load, with the rationale that viral infection could either directly impact ovary function by infecting the tissue, or indirectly impact ovaries by shunting resources into immune activation while depleting resources available to invest in ovarioles.

We first tested zero-inflated statistical models using total viral load (DWV, SBV, and BQCV copies combined; **Figure 1a**) as a continuous variable, but all models suffered from residual overdispersion; therefore, we binned the viral data into the categories “low” (0 copies), “medium” (0 – 10^5^ copies), and “high” (10^5^ < copies) (**Figure 2b**) and used a linear mixed model. Among the n = 83 queens for which both viral data and complete fertility metrics were acquired, we found that the total viral load was significantly linked to ovary mass (**Figure 2c**; low-high contrast: z = -2.24, df = 71, p = 0. 028; medium-high contrast: z = -1.35, df = 71, p = 0.18). There was neither a significant relationship between ovary mass and sperm counts nor sperm viability; therefore, these parameters were dropped from the final statistical model. See **Table 2** for complete statistical reporting and **Supplementary Table S3** for the underlying data. The significant relationship between ovary mass and viral load is not sensitive to the exact cut-offs for binning the viral data; in fact, an even greater significant effect is observed if the cut-off between “medium” and “high” bins is set to 10^4^ copies instead of 10^5^ copies (low-high contrast: z = 2.87, p = 0.0051).

**Figure 2.**
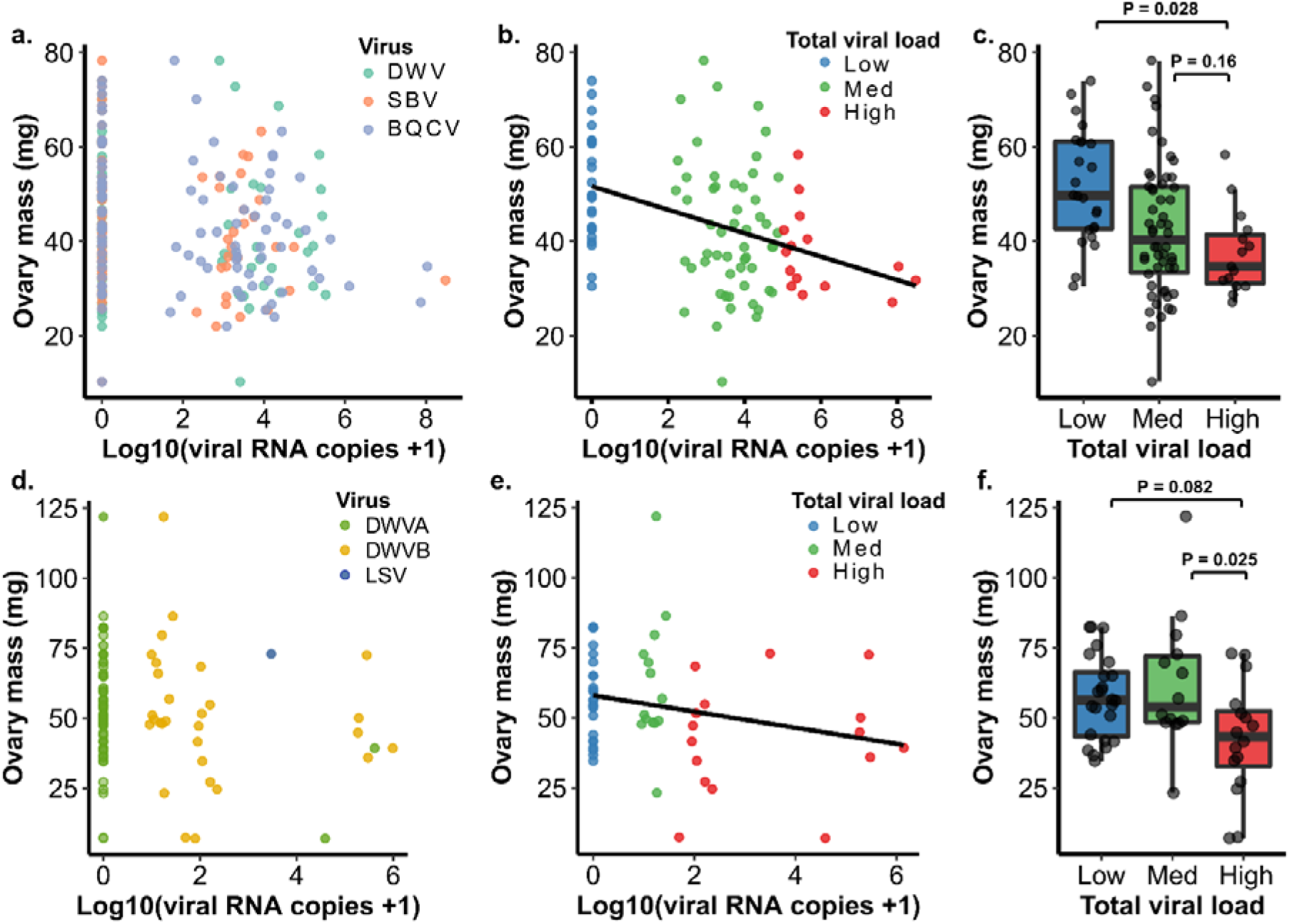
Relationships between viral RNA copies and ovary mass. See **Table 1** for complete statistical details. A) Survey 1 (previously published; see Table 1 for details) included viral RNA count data for deformed-wing virus (DWV), sacbrood virus (SBV), and black queen cell virus (BQCV), using head tissue samples. B) Viral RNA copies for each virus were added before transformation to produce the variable “Total viral load,” which we split into three categories to simplify statistical analysis. C) Ovary mass was modeled against queen health status (healthy or failed), sperm viability, sperm counts, and total viral load using a linear mixed model, with queen source (apiary) as a random effect. Sperm viability and sperm count variables were not significant and were dropped from the final model. D) We obtained a validation data set (Survey 3) of n = 54 queens with known ages. We analyzed 8 different viruses in the thorax using a different laboratory service. Queens were mainly infected with DWV-B (non-detected viruses not shown), with sporadic DWV-A and Lake Sinai virus (LSV). E) Viral copies were combined into a total viral load as in b). Different cut-offs were necessary owing to dramatically different copy distributions compared to b). f) We validated the negative relationship between ovary size and total viral counts. We used a linear mixed model with queen age (0, 1, or 2 y old), health status (healthy or failed) and total viral load (low, medium, and high) as fixed effects and queen source (apiary) as a random effect.

We further validated this observation with the n = 54 queens from Survey 3, which we sampled independently from different apiaries, and whose viral abundances were analyzed by a different laboratory (data in **Supplementary Table S4; Figure 2d-e**). In this data set, we were also able to account for queen age in our statistical model, which ranged from 0 to 2 y for both failed and healthy queens. Despite identifying an overall lower range of viral abundance (0 – 10^6^ copies compared to 0 – 10^8^ copies, likely owing to different laboratory methods), the overall negative relationship between viral titer and ovary mass remained significant (low-high contrast: z = -1.78, df = 43, p = 0.082; med-high contrast: z = - 2.32, df = 43, p = 0.025; **Figure 2f**), demonstrating that this is a reproducible result. The different range in total viral copies identified in this dataset necessitated different binning cut-offs (“low”: 0 copies; “medium”: 0 – 10^1.5^ copies; “high”: 10^1.5^ < copies) than Survey 1 data (“low”: 0 copies; “medium”: 0 – 10^5^ copies; and “high”: 10^5^ < copies). If the same bin definitions are used the relationship between ovary mass and viral load is no longer significant, likely as a result of only five queens qualifying for the “high” category.

Temperature stress and pesticide stress are factors which have been previously associated with fertility declines^37,42-45^, but despite our previous efforts^38^ we currently have no way of quantifying the exposure of the queens in our field surveys^46,47^. Therefore, these are potential extraneous variables which could influence our results. To determine if these stressors could have a confounding effect in our data, we tested the effect of heat stress and pesticide exposure separately and found no effect on ovary mass (statistics are reported in **Table 2**; see **Supplementary File S1 and Supplementary Tables S5 and S6** for underlying data). We thus rule them out as influential factors, strengthening our confidence in viral titers being the main driver for ovary mass declines as depicted in **Figure 2**.

To confirm that viral infection is a cause of the reduced ovary size associated with queen failure and not merely a result of failed queens potentially being more susceptible to infection, we injected age-matched, mated queens with IAPV. We chose IAPV because it naturally infects honey bee queens, but has a low frequency of infection^39^, so experimental infection is unlikely to confound with natural abundance. After just 65 hours, we found that the IAPV-injected queens had significantly smaller ovaries relative to the sham-injected queens (t = -3.185, p = 0.005) (**Figure 3a**), echoing our results from the field.

**Figure 3.**
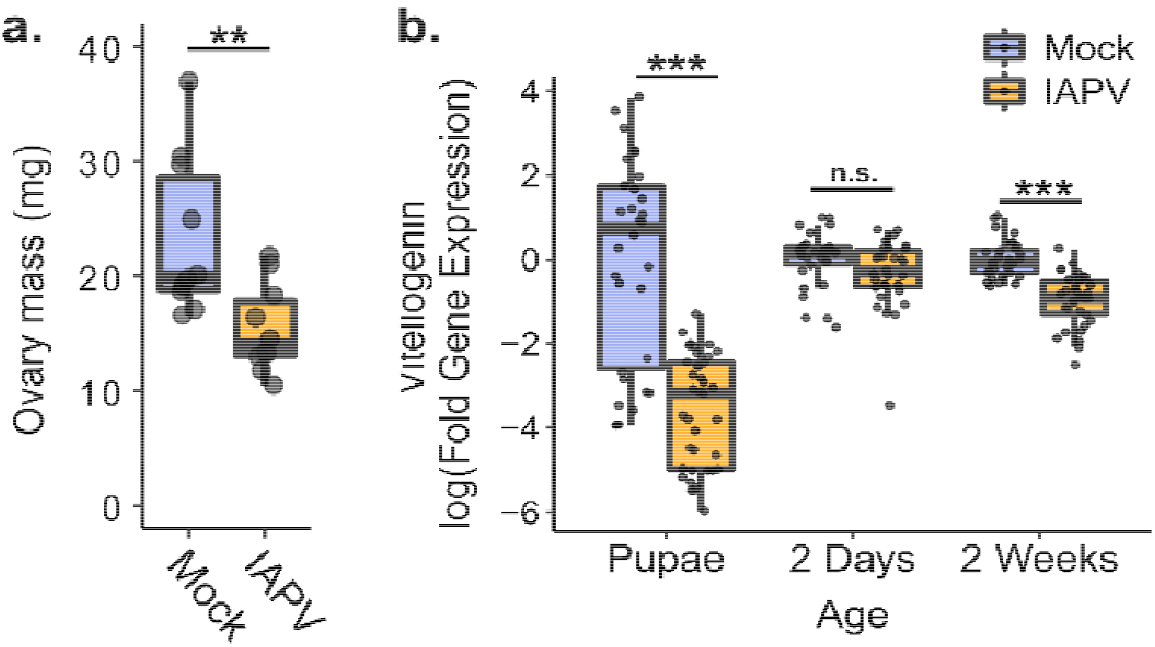
Ovary mass and vitellogenin transcription are significantly decreased after IAPV injection. a) Mated, age-matched queens were injected with IAPV or a sham (buffer only; n = 10 each). After 65 hours the ovary mass of infected queens was significantly reduced. Significance was evaluated using a t test. ** p = 0.005. b) The transcript for vitellogenin is downregulated in the abdomens of queens experimentally infected with IAPV at two days before emergence and two weeks post-emergence (after CO_2_ ovary activation). Fold gene expression was calculated using the 2^−ΔΔCt^ method. Pairwise comparisons of gene expression were evaluated using the Wilcoxon Rank Sum test (n.s. = not significant, *** = p < 0.0001).

In a separate experiment, we investigated vitellogenin gene expression in the abdomens of IAPV-injected queens relative to sham-injected queens at three different ages: two days before emergence (pupae), two days after emergence, and two weeks after emergence (with CO_2_ treatment to stimulate ovary activation and egg laying^48^). The success of the infections was confirmed by measuring IAPV copies using qPCR (**Supplementary File 1**.) In pupal queens and queens two weeks after emergence (after CO_2_ ovarian stimulation), we found that expression of vitellogenin was significantly reduced by IAPV infection (**Figure 3b**). Vitellogenin, the major egg yolk protein precursor, is one of the most significant sources of nutrition for developing oocytes, where is it deposited in massive quantities. Much of the mass of a queen’s ovary is made up of developing eggs, therefore a decrease in vitellogenin expression is consistent with a reduction in ovary mass and together represents a reduction in the resources being invested into producing or provisioning eggs. Vitellogenin is also responsible for the binding and transport of immune elicitors (*i*.*e*., bacterial cell wall components) during transgenerational immune priming^49,50^. So, this reduction in vitellogenin could have knock-on effects for the success of the queen’s progeny by reducing her capacity to provide them with immune priming.

### Ovaries of failed queens have altered protein composition

To determine if the changes we observed in the ovaries of failing queens were associated with altered patterns of protein expression consistent with a disruption in egg production, we performed quantitative proteomics on the ovaries of n = 88 queens collected in British Columbia from Survey 1. We found that 415 proteins, around 20% of the total number of quantified proteins, were differentially expressed between failing and healthy queens at a false discovery rate of 5% (Benjamini Hochberg method; **Figure 4a**). Many of the proteins that were downregulated in failing queens are associated with metabolic processes, but no gene ontology (GO) terms were significantly enriched among the differentially expressed proteins. This could be due to the poor GO characterization of many honey bee proteins, as we have previously described^36^. However, among these differentially expressed proteins are several indicators of an increase in immune activity and decrease in egg development (**Figure 4b**). One of the top proteins decreased in failed queens was prohibitin (XP_624330.3), a highly multifunctional protein associated with cellular proliferation^51^, the regulation of follicular development in mammalian ovaries^52^, and a mediator of viral entry into insect cells^53^. Furthermore, heat-shock proteins have antiviral activity in insects^54,55^, and our data are consistent with that role in queens. We found that two small heat shock proteins (protein lethal(2) essential for life), XP_001119884.1 and XP_001120194.1, as well as dnaJ homolog shv (XP_006569897.2) were upregulated in failed queens. Despite sharing the same name (protein lethal(2) essential for life), these proteins do not share tryptic peptides > 6 residues long; therefore, their quantification was not influenced by shared peptide sequences. DnaJ homolog shv was also previously reported to be associated with the antiviral heat-shock response in workers^55^.

**Figure 4.**
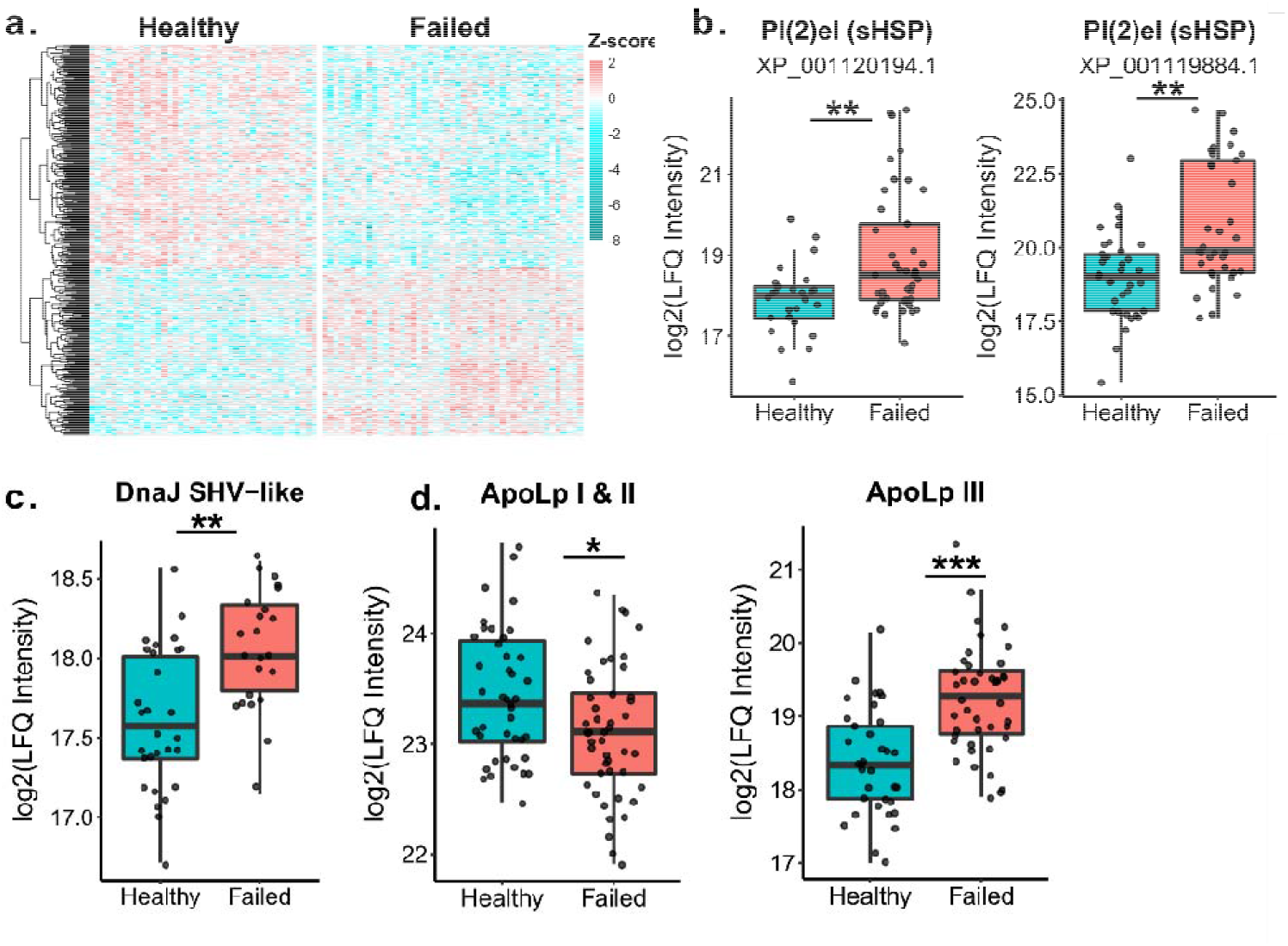
Changes in ovary protein expression and vitellogenin. Protein abundance was measured via LC-MS/MS using label-free quantitation (LFQ). Analysis was done on proteins identified in at least 10 samples and expression is reported as log2-transformed LFQ intensity. We used limma with status (failed vs. healthy), ovary mass, and total viral counts as fixed effects for n = 88 queens (41 healthy and 47 failed) to identify the differentially expressed proteins shown in a, b, and c. Exact sample sizes may differ due to missing values for some proteins. The false-discovery rate (FDR) was controlled using the Benjamini-Hochberg method (5% FDR). a) Protein expression patterns in ovaries of healthy and failed queens. Only proteins differentially expressed and quantified in 75% of samples are shown (387 proteins). b) Proteins associated with the antiviral response are upregulated in failed queens (sHSP (XP_001119884.1): t = 3.8, adjusted p = 0.003; sHSP (XP_001120194.1): t = 3.3, adjusted p = 0.01; DnaJ: t = 4.9, adjusted p = 0.0003). c) Apolipophorins I/II are involved in lipid transport and are downregulated in failed queens (t = -2.59, adjusted p = 0.048). Apolipophorin III is an important protein for immune function and lipid transport and is upregulated in failed queens (t = 5.25, adjusted p < 0.0001).

Studies in Lepidopterans and Orthopterans have shown that the protein apolipophorin III (apoLp-III) is involved in both the immune system and metabolism^56^. When in its monomeric form, it acts as an immune surveillance pattern-recognition receptor with the ability to activate the immune response. We previously found that apoLp-III is significantly co-expressed with a cluster of predominantly innate immune proteins in the spermathecal fluid^36^, lending confidence to its involvement in immune processes in queens. But, when combined with high-density apolipophorin (apoLp-I/II), it acts as a lipid transporter, shuttling lipids liberated from the fat body and from the digestive system around the body^57^. One of the most significant destinations of these lipids in reproductively active female insects are the developing oocytes, where they will make up 30-40% of each egg’s dry weight^58^. These lipophorins could be a molecular switch governing both the movement of resources and immune function, and we predicted that if increased viral loads are a feature of failed queens, then they should have increased levels of apoLp-III to promote immune function. Additionally, we predicted that apoLp-I/II, which is constitutively involved in lipid transport and juvenile hormone signalling, would be reduced in the smaller ovaries of failing queens which are likely producing fewer eggs. We found that apoLp-III was indeed upregulated in failed queens, and apoLp-I/II was downregulated (*Figure 4c*), suggesting that lipid transport to developing eggs is reduced in failing queens.

### Heat-shock proteins are upregulated in virus-infected queens

To gauge whether the levels of viral infection in the field were sufficient for the queens to launch an antiviral response, we investigated expression of putative antiviral proteins. Virus-infected workers express elevated levels of antimicrobial peptides (*e*.*g*., *defensin, hymenoptaecin*); however, these peptides’ molecular properties are not favourable for quantitation by shot-gun proteomics without specialized enrichment strategies. Previous work has demonstrated that in fruit flies and worker honey bees, heat-shock proteins, which are more easily quantified by proteomics, are an important part of the antiviral defense^54,55,59^. Virus-infected workers also upregulate heat-shock protein mRNA (*pl(2)el, hsp70-3, hsp70-4, hsp83-like*, and *hsp90*), and workers whose heat-shock proteins were experimentally induced by temperature stress shortly after infection had 74-90% lower viral titers compared to non-induced controls^55^. We therefore hypothesize that antiviral heat-shock protein expression is also correlated with viral load in queens.

We first analyzed previously published quantitative proteomics data obtained from spermathecal fluid samples of the queens from Survey 1 with varying degrees of viral infection^36,38^. Honey bee viruses can be transmitted sexually, so spermathecal fluid is a reasonable tissue in which to look for an antiviral response^60^. Our statistical model included sperm counts, health status, and log-transformed total viral RNA copies as fixed effects. We found that six proteins significantly correlated with log-transformed total viral RNA copy numbers at a false discovery rate of 10% (**Table 3**), half of which are heat-shock proteins: heat-shock protein 70 cognate 4 (HSP70-4) and two other small heat-shock proteins (sHSPs) XP_001120006.2 and XP_001119884.1. As we predicted, all three HSPs positively correlate with viral abundance (HSP70-4 and sHSP XP_001120006.2, which have been previously associated with the worker bee antiviral response^55^, are depicted in **Figure 3a-b**). The other three proteins—a transmembrane protease, zinc carboxypeptidase, and arylsulfatase-B—have not been previously linked to viral infection in honey bees. Notably, the two sHSPs were previously proposed as candidate diagnostic markers for heat-stress and were significantly upregulated in failed queens relative to healthy queens^38^; here, we show that they are significantly correlated with viral abundance even when health status is included as a fixed effect. This observation unfortunately negates their utility as heat stress biomarkers due to confounding virus infection.

**Table 3.** Statistical summaries of proteins correlating with total viral loads.

| Protein | Common name | \|t\| | P | Adjusted p* |
| --- | --- | --- | --- | --- |
| XP_016771183.1 | Transmembrane protease serine 11B like | 4.48 | 0.0000355 | 0.054 |
| XP_001120006.2 | Protein lethal(2) essential for life (small HSP) | 4.17 | 0.0000778 | 0.054 |
| NP_001153522.1 | Heat shock protein 70 cognate 4 | 4.04 | 0.000114 | 0.054 |
| XP_623922.2 | Zinc carboxypeptidase | 4.05 | 0.000136 | 0.054 |
| XP_006569022.1 | Arylsulfatase-B | 4.81 | 0.000101 | 0.054 |
| XP_001119884.1 | Protein lethal(2) essential for life (small HSP) | 3.82 | 0.000246 | 0.082 |

\*Benjamini Hochberg method

To validate that these HSPs are part of the antiviral immune defense in queens we performed experimental infections with IAPV. We selected the two heat shock proteins that we identified as upregulated in naturally virus infected queens that overlapped with the previously reported list of HSPs associated with the antiviral response in workers: protein lethal(2) essential for life (XP_001120006.2, Pl(2)el) and heat-shock protein 70 cognate 4 (Hsp70-4)^55^ (all primer sequences are available in **Supplementary Table S7**). Pl(2)el and HSP70-4 were significantly upregulated after viral infection in queens of all three ages, with the exception of two-week-old queens, in which HSP70-4 was not differentially expressed (**Figure 3c-d, Supplementary Table S8**). The fact that HSP70-4 was not upregulated in queens whose ovaries had been stimulated to develop may be because insects which have mated or begun to reproduce often demonstrate a reduced ability to mount an immune response relative to their unmated counterparts^27^, and trade-offs involving the immune response often involve a complicated reconfiguration in which different components of the response might become up or downregulated^61^.

**Figure 3.**
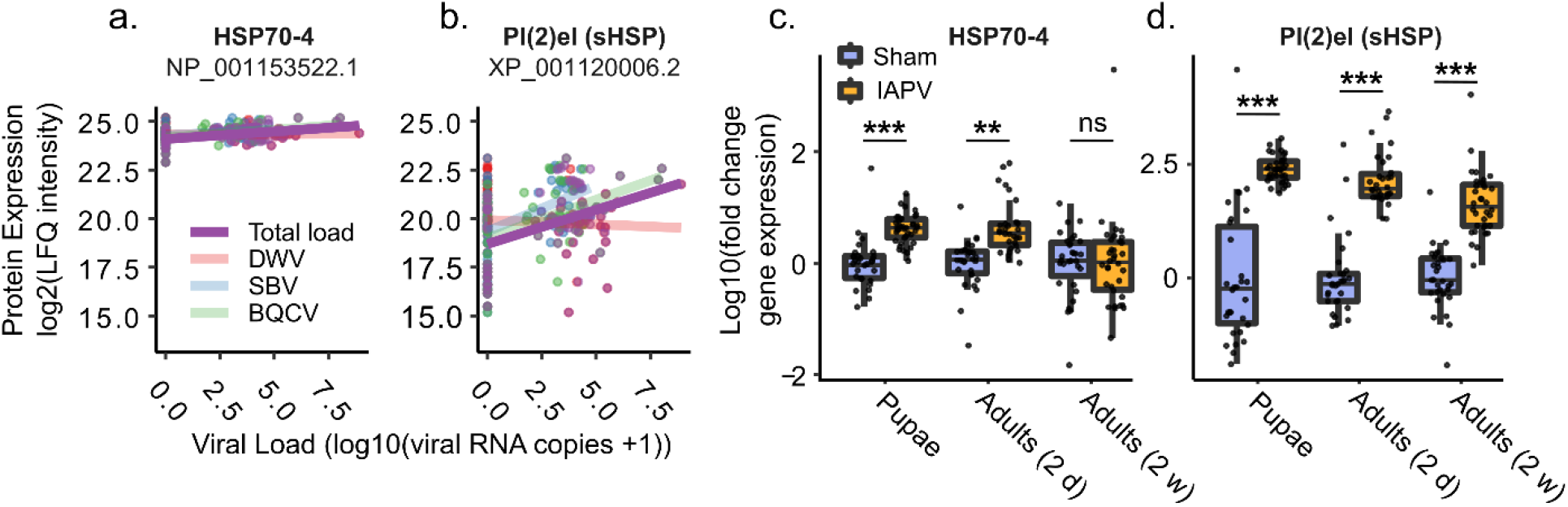
Heat-shock proteins associated with natural and experimental viral infections. a & b) We analyze previously published proteomics data using limma, including health status, sperm counts, and log transformed total viral loads as fixed effects (n = 88 queens had complete proteomics with no missing values in continuous covariates). Individual virus types are shown, but only total titers were statistically analyzed. P values were corrected for multiple hypothesis testing by Benjamini-Hochberg correction (see **Table 3** for statistical summaries of these and other significant proteins). c & d) The transcripts of two heat-shock proteins found in the spermathecal fluid of naturally infected queens and previously identified in the antiviral response in workers are upregulated in the abdomens of queens experimentally infected with IAPV. Fold gene expression was calculated using the 2^−ΔΔCt^ method. Pairwise comparisons of gene expression were evaluated using the Wilcoxon Rank Sum test. ns: not significant; ** = p < 10^−7^; *** = p < 10^−10^.

## Discussion

Here, we show through experimental injections that virus infection shrinks queens’ ovaries, reduces vitellogenin expression, and stimulates the expression of putative antiviral immune effectors (HSPs). The observation that queens deemed to be “failing” by beekeepers had higher viral loads and smaller ovaries than queens perceived to be “healthy” suggests that this is an economically relevant phenomenon. Furthermore, using a different virus from those which we identified in the queens from the field provides evidence that this upregulation of the antiviral response and reduction in reproductive activity are general responses to viral immune activation, and not a specific response to a particular virus. The negative impact of viral infections could either be due to direct impacts of the viral pathology, or indirect impacts of forcing the queens to split resources between reproduction and launching an immune defense. Regardless of the mechanism, the data show that a high level of natural viral infection in field-sampled queens is linked to small ovary size and reduced reproductive output, underscoring the likelihood that viral infections are having a biological impact on queens in the field.

Honey bees and other highly social insects, such as ants, have escaped the trade-off between reproduction and lifespan, which is present in most insects^34,62-64^. Highly reproductive individuals in other species are typically short-lived, but Hymenopteran and termite queens exhibit both enormous reproductive output and extraordinary longevity compared to the non-reproductive castes. While surviving infection is clearly tied to being long-lived, the extent to which honey bees and other social insects may have escaped the specific trade-off with immunity is yet unclear. Recent work has shown that reproductive activation of honey bee workers protects them against oxidative stress and improves survival after virus infection^65^, and these results suggest a potential escape from the reproduction-immunity trade-off. Furthermore, some research suggests that queens of a black garden ant, *Lasius niger*, have also escaped the reproduction-immunity trade-off when stimulated with a fungal pathogen^66^, but responses of other species to other pathogens, including viruses, may differ.

The ovaries are the most likely tissue that may manifest a reproduction-immunity trade-off due to compromised resource allocation because they are the biggest resource sink related to reproduction^35^. Laying their body weight in eggs per day is a massive investment of resources in a singular reproductive process, contrary to sperm maintenance, which is achieved mainly by enzymatic suppression of reactive oxygen species in the spermatheca and creating the conditions that sustain sperm in a metabolically quiescent state^67,68^. Indeed, if reproduction-immunity trade-offs apply to sperm storage, as has been suggested in honey bees and other insects^27,32,33,36,69^, they would most likely be due to collateral damage of immune effectors, such as reactive oxygen species, rather than resource-allocation.

Indeed, ovary mass tends to decline when queens are confined to small cages with only a few attendants during shipping, but it rebounds again after being placed in a queen bank colony for several days or when she reinitiates oviposition^38^. Therefore, the effects that we observe on ovary size and protein composition may be temporary and could conceivably be reversed if, for instance, the virus infection was cleared. Our findings of an upregulation of antiviral effectors associated with a decrease in indicators of ovary activity are consistent with previously published findings involving the reproductive-immunity trade-off in other insects^29,70,71^. However, we cannot rule out the possibility that the effects we observe on the ovaries are some direct pathogenic result of viral infection.

In conclusion, our data show a clear relationship between ovary size and viral infection. Small ovary size and viral abundance are also associated with queen failure, a finding that we validated with an independent sample of queens. Protein expression patterns in the ovaries of failed queens indicate an antiviral response and a concomitant decrease in oocyte development. Furthermore, we show that additional stressors (heat and pesticide stress) with unknown prevalence in our sampled queen population are not associated with reduced ovary mass. The increase in expression of antiviral HSPs we observed in naturally infected queens was recapitulated in queens of different ages experimentally infected with a different virus. Since some of the same antiviral proteins are upregulated with both virus infection and heat-shock, these data also highlight a potential mechanism by which extreme temperatures associated with climate change could have immunomodulatory effects. Whether such an interaction would have positive or negative impacts on reproduction remains to be tested. Regardless, the present data suggest that virus-specific effects are driving ovary mass reductions. It is possible that pathogen spillover events, as has been suggested to occur between honey bees and other species, could negatively impact insects not only through direct fitness costs of infection, but indirect effects on the population via reduced reproductive output.

## Materials and Methods

### Queen surveys

For Survey 1, we utilized previously published data^36^, which included N = 105 queens (excluding imported queens, which were biased to smaller ovary sizes due to a longer caging duration). N = 93 of these queens had associated viral data. For the ovary mass comparison across surveys, we retained only queens from producers that met the following criteria: the producer collected both failed and healthy queens, queens were delivered to the laboratory in one shipment, and the producer provided n > 2 failed and healthy queens each. These criteria were applied because in the 2019 survey, producers from many different locations shipped queens to the laboratory by different methods and with collection schedules; therefore, queens spent different lengths of time in transit and in cages prior to transit, which we know from previous work impacts ovary mass. Applying these criteria reduced our sample size for the ovary mass comparison but removed these extraneous effects.

For Survey 2, N = 22 queens were collected from colonies in 2018 from three different apiaries in southeastern Pennsylvania. Healthy and failed queens were age- and apiary-matched in order to eliminate sampling bias. The queens were frozen on dry ice immediately in the field and shipped frozen to the University of British Columbia, at which time the queens were thawed on ice and dissected.

For Survey 3, queens were collected from local collaborating beekeepers as they became available during the summer of 2020. Apiary location, colony symptoms (if failed), genetic background (*i*.*e*., local or imported stock), and age were recorded. The beekeepers considered the queens to have ‘failed’ if they either displayed spotty brood patterns, were drone layers, or if the adult population rapidly dwindled, including the presence of chalkbrood infections that the colony could not resolve on their own. Beekeepers considered the queens to be ‘healthy’ if they produced robust brood patterns, the colony did not have signs of appreciable brood infections, and the adult population was strong. Failed and healthy queens were caged with candy and attendants then kept in a queen bank with plentiful nurse bees if they could not be immediately analyzed in the laboratory. The queens were transported to the University of British Columbia, where they were anesthetized with carbon dioxide, decapitated, and their spermathecae were removed for sperm viability analysis. The remaining queen bodies were stored at -70°C until further dissection to determine ovary mass.

### Sperm viability, sperm counts, and ovary mass

Sperm viability measurements for Survey 3 queens were obtained essentially as previously described^36^, following the methods of Collins and Donaghue^72^. Briefly, we burst the spermatheca with tweezers in 100 μl of Buffer D (17⍰mM D-glucose, 54⍰mM KCl, 25⍰mM NaHCO_3_, 83⍰mM Na_3_C_6_H_5_O_7_) and gently agitated the solution until sperm cells were homogeneously dispersed. We then stained the sperm using SYBR-14/propidium iodide dual fluorescent staining following the manufacturer’s protocol (Live/Dead sperm viability kit, Thermo). We dispensed 5 μl of the stained sperm into a well of a 24-well plate, over which we placed a round glass cover slip. We imaged the sperm (three fields of view per queen, average of ∼100 sperm cells per image) using a Cellomics ArrayScan XTI (Thermo) and automatically counted live (green) and dead (red) channels using ImageJ v1.52a. Relative sperm abundances were obtained by counting the cumulative number of sperm across three fields of view. Every sample was analyzed in the same style of plate and with the same sized cover slip, so measuring the total sperm in a given area is proportional to total sperm in the sample. We later dissected the ovaries from the queen remains and weighed them using an analytical balance. Data for Survey 1 is already published and we direct the reader to McAfee *et al*.^36^ for the methods regarding the collection and processing of those queens.

### Viral quantification for queen surveys

Viral data (SBV, DWV, and BQCV) for Survey 1 were obtained by RT-qPCR exactly as previously described and are previously published data^36^. Briefly, samples were submitted to the National Bee Diagnostics Center, where total RNA was extracted from the head using a guanidinium isothiocyanate extraction buffer, cDNA was synthesized using the iScript cDNA synthesis kit (Bio-Rad Laboratories, Hercules, USA), and virus RNA was quantified by qPCR with previously published primers^36^. Standard curves for each virus amplicon were generated using serially diluted plasmids containing the target sequence. RP49 was used as the honey bee reference gene, enabling the viral RNA copy number per queen head to be calculated. Copy numbers per head of all three viruses were added together to generate the ‘Total counts’ variable. Queens from Survey 2 were not analyzed for viral infections because the sample size (22 queens) is too small to yield sufficient statistical power linking viral abundance to ovary size.

Queens from Survey 3 were processed by the Honey Bee Queen and Disease Clinic at North Carolina State University using methods as previously described^73^ with minor adjustments. For this sample set, we analyzed thoraxes because we have observed that other components of the head can sometimes interfere with PCR. We also tested for a wider range of viruses, including DWV-A, DWV-B, ABPV, SBV, IAPV, LSV, BQCV, and CBPV. Thoraxes were pulverized by bead beating and total RNA was extracted via the Zymo Research DirectZol RNA miniprep kit and Trizol reagent. RNA was quantified and verified for quality by NanoDrop Spectrophotometry and diluted to 200 ng/μL in RNAse/DNAse free water. cDNA was synthesized using the BioBasic All-In-One RT MasterMix. qPCR was performed on a QuantStudio 5 thermocycler using PowerUp Sybr Green chemistry. Standard curves were generated using serial dilutions of known quantities of custom plasmids containing the target sequences. Actin, 28s ribosomal subunit, and GapDH were used as reference genes. Final copy numbers were normalized to reference gene values using GeNorm software^74^.

### Queen stress experiments

Italian queens were imported from California for the pesticide stress experiment, whereas the queens for temperature stress were grafted from existing colonies, reared using standard techniques^75^ at the University of British Columbia and mated locally. The temperature stress experiments were conducted in June 2019 and the pesticide stress experiments were conducted in August 2019. Queens were introduced to established 3-frame nucleus colonies and allowed to freely lay eggs for 2 weeks prior to exposing them to stressors. Colonies were fed protein patties (Global) and sugar syrup for the duration of the experiment. Prior to stressing, queens were caged with six attendant workers and candy, transported to the laboratory, anesthetized with carbon dioxide (5 min), and weighed on an analytical balance.

Once revived, queens were temperature stressed by placing them and their attendants in a 42°C incubator, humidified with a water pan, for 2 h. Control queens were placed in a 33°C incubator for 2 h. For the pesticide stress experiment, queens were administered 2 μl of a pesticide cocktail, diluted in acetone to a hazard quotient of 3,500, directly on the thorax while still comatose. We have previously used this pesticide cocktail at a dose of 2 μl diluted to a hazard quotient of 511 in order to investigate queen detoxification response^38^, but here we used a hazard quotient of 3,500 because this is the approximate contamination level in wax that has been previously associated with ‘queen events’^42^. The cocktail is composed of coumaphos, tau-fluvalinate, 2,4-dimethylphenyl formamide (an Amitraz degradate), chlorothalonil, chloropyrifos, fenpropathrin, pendimethlalin, atrazine, and azoxystrobin mixed in the same relative proportions as McAfee et al^38^. Solvent control queens received acetone alone, and handling controls received no treatment.

The stressed queens were then transported back to the apiary and reintroduced to their respective hives (2 d re-acclimation, then release). They were allowed to freely lay until two weeks after the stress event, then they were again caged and transported to the laboratory for euthanasia and dissection. Fertility metrics were measured as described above.

### Statistical analyses of phenotypic data

All statistical analyses were performed in R v3.5.1. For Survey 3 data, we first checked if ovary mass, sperm count, and sperm viability were normally distributed and had equal variance between failed and healthy queens using the Shapiro and Levene test, respectively. The ovary mass and sperm viability passed these tests (p > 0.05); therefore, a linear mixed model (lme) was used for both, including health status (failed vs. healthy) and age (0, 1, or 2 years) as fixed effects and location (9 levels) as a random effect. The sperm viability data, however, was not normally distributed, so we analyzed these data with a generalized linear mixed model with a logit link function (glmer, family = binomial) as previously described^33^, constructed using the same fixed and random factors, but where sperm viability was represented as counts of ‘successes’ (live) and ‘failures’ (dead). Ovary mass data from three independent queen surveys were analyzed using a linear mixed model (lme) with health status as a fixed effect and queen source location as a random effect.

To determine the link between ovary mass and virus infection, we first tested used total viral loads as raw counts and tested the appropriateness of zero-inflated regression models using a Poisson distribution and negative binomial distribution but found that residuals suffered from over dispersion (dispersion parameter >> 1). We repeated these tests using log2 transformed viral data, but residuals were still over-dispersed. Therefore, we binned total viral loads into “low” (virus not detected), “medium” (0 < log10(total virus +1) ≤ 5), and “high” (log10(total virus +1) ≥ 5) infection groups to simplify the statistical analysis. This yielded n = 31, 56, and 19 queens in each group, respectively. Ovary mass was modeled against queen health status (healthy or failed), sperm viability, sperm counts, and total viral load using a linear mixed model, with queen source (apiary) as a random effect. Sperm viability and sperm count variables were not significant and were dropped from the final model. Total viral counts of Survey 3 queens were also binned into three infection categories, but with different cut-offs owing to different abundance distribution (likely owing to samples being analyzed by a different lab and from a different tissue). Survey 1 virus data was obtained from the head, whereas Survey 3 viral data was obtained from the thorax. For Survey 3, the bins were: low (virus not detected), medium (0 < log10(total virus +1) ≤ 1.5), and high (log10(total virus +1) ≥ 1.5), yielding n = 24, 14, and 16 queens. In the higher range of viral loads, the data are broadly dispersed (see **Figure 2e**) and setting a higher threshold for the “high” bin yielded too few queens in this category. We analyzed the data with a linear mixed model with queen age (0, 1, or 2 y old), health status (healthy or failed) and total viral load (low, medium, and high) as fixed effects and queen source (apiary) as a random effect.

For the comparison of weights and ovary masses of queens exposed to either heat stress or pesticide stress, we first checked for data normality and equal variance as described above. The data passed these tests (p > 0.05); therefore, we performed statistical analysis on pesticide-stressed and heat-stressed queens separately using a simple linear model (lm) with treatment group as a fixed effect. For the short-term heat stress experiments, ovary masses were first mean-centered by experiment, then analyzed using a simple linear model with treatment as a fixed effect.

### Ovary proteomic sample preparation and statistical analysis

Ovary samples from Survey 1 (N = 88, 5 samples from the original 93 queens were lost during processing) were prepared for mass spectrometry essentially as previously described^37^. One ovary from each queen was homogenized in a separate 2-ml screw cap tube containing 300 μl of lysis buffer (6 M guanidinium chloride, 100 mM Tris, pH 8.5) and four ceramic beads. The homogenizer (Precellys 24, Bertin Instruments) was set to 6,500 s^-1^ for 30 s: samples were homogenized three times and placed on ice for 1 minute between each. Then samples were centrifuged (16,000 relative centrifugal force, 10 min, 4 °C) to remove debris. 100 μL of supernatant was transferred to a new tube and diluted 1:1 with distilled H_2_O. Protein was precipitated by adding four volumes of ice-cold acetone and incubating overnight at −20 °C. The precipitated protein was pelleted by centrifugation (6,000 relative centrifugal force, 15 min, 4 °C) and the supernatant was discarded. The protein pellet was washed twice with 500 μl of 80% acetone, then the pellet was allowed to air dry (∼5 min) before solubilization in 100 μl of digestion buffer (6 M urea, 2 M thiourea). Approximately 20 μg of protein was reduced (0.4 μg dithiothreitol, 20 min), alkylated (2 μg iodoacetamide, 30 min, dark), and digested (0.4 μg Lys-C for 2.5 h, then 0.5 μg trypsin overnight). Digested peptides were acidified with one volume of 1% trifluoroacetic acid and desalted with high-capacity STAGE tips as previously described^76^. Eluted samples were dried (SpeedVac, Eppendorf, 45 min) and resuspended in Buffer A (0.1% formic acid, 2% acetonitrile), then peptide concentrations were determined using a NanoDrop (Thermo, 280 nm). LC–MS/MS data acquisition was performed as previously described^37^.

MS data was searched using MaxQuant (v1.6.8.0) with the parameters and database as previously described^37^. In Perseus (v 1.5.5.3), the resulting protein groups were filtered to remove reverse hits, contaminants, proteins identified only by site, and proteins identified in fewer than 10 samples. Normalized LFQ intensity was then log2 transformed. Subsequent analyses were performed in R (v3.6.3). Differential expression analysis was performed using the limma package in R^77^ with ovary mass, status (failed vs. healthy), and total viral counts (log10 transformed of combined RT-qPCR counts, +1 to avoid undefined terms) included as fixed effects. Empirical Bayes moderation of the standard errors was then performed on the resulting linear model, and finally the false discovery rate was controlled to 5% (for the status coefficient) or 10% (for the ovary mass coefficient) using the Benjamini-Hochberg correction. GO term enrichment analysis was performed on the proteins differentially expressed among status groups using Ermine J as previously described^38^, though no terms were significant. We used a linear model with ovary mass, status, and total viral counts as fixed affects to analyze the expression of ApoLp-I/II and ApoLp-III.

### Statistical analysis of spermathecal proteomic data

We analyzed previously published proteomics data using the limma R package essentially as previously described for ovaries. Proteomics data were first groomed to remove samples that did not have associated viral abundance data. We included log10 transformed total viral load (+1 to avoid undefined terms), health status (failed or healthy), and sperm counts as fixed effects in the statistical model. We used a Benjamini-Hochberg correction to control the false discovery rate to 10%. We direct the reader to McAfee *et al*. for details on proteomics sample preparation and upstream data processing.

### IAPV experimental infections for measuring ovary mass

Mated queens of the same age were obtained in a single shipment from California. Queens were anesthetized with CO_2_ for 6 minutes before being injected between the abdominal tergites with approximately 4nL of IAPV-inoculum (n=10) or phosphate buffer saline (PBS) as sham (n=10). Injections were performed using the FemoJet Microinjector (Eppendorf) with hand-pulled glass needles. The queens were then kept in wooden cages with 5 attendant workers and provided with sugar candy in the incubator at 34°C with a water pan for humidity. After 65 hours the queens were sacrificed, and the wet ovary masses measured on an analytical balance. The dilution of purified IAPV used to inject the queens was determined after measuring survivorship in workers injected with a dilution series. In this experiment, attendants in the cages with IAPV-injected queens were beginning to show symptoms of paralysis after 2.5 days (presumably from contact with the infected queens).

### IAPV experimental infections for expression analysis

Three large groups of sister queens were produced from a healthy mother queen to study the direct effect of virus infection on queens of three different ages following honey bee queen-rearing technique^75^. Briefly, young larvae (12 – 24 hours old) were grafted into the artificial queen cups and placed in a populous but queen-less nurse colony to develop. Prior to grafting, the donor colony was visually determined to be free of symptomatic diseases (Nosemosis, Varroasis, European and American Foulbrood, and Chalkbrood). After 6 days, sealed queen cells were transferred to an incubator (35 °C, 65% humidity) and treated as follow:

The first group was used for pupal infections. Two days before emergence, queen pupae were gently removed from their capped cells. These pupae were either injected with 2 μL of IAPV-inoculum containing 5.24×10^3^ virus particles (n= 34), or Phosphate buffered Saline (PBS) as a sham control (n= 31). The injection was performed using a Nanojet™ syringe pump (Chemix, USA) with an infusion flow rate of 0.1 μL/s, where the needle was inserted into the first abdominal segment located immediately behind the metathorax. Each queen then was placed individually in a 15 mm well in the 24 well glass bottom cell culture plate and kept in the incubator (35 °C, 65% humidity). While in the incubator, development of the virus injected queens was compared with the PBS injected queens. Our observation confirmed a complete cessation of development in IAPV injected queens 20 hours post injection, not observed in the control group. Therefore, the experiment was terminated, and the abdomen of each queen was then separated, placed in an Eppendorf tube, and stored in a freezer at -80 °C until RNA extraction.

The second group of queens was used for infecting newly emerged queen in the pre-reproductive stage. Two days after emergence, queens were either injected with the IAPV-inoculum (n= 30), or PBS as shame control (n= 30) as explained above. Each queen was accompanied with 10-15 worker honey bees from the donor colony in a single-use plastic cup, provided with water and sugar candy. Cups of IAPV-inoculated and PBS-injected groups were maintained separately in the incubator (35 °C, 65% humidity).

The IAPV-inoculated queens started showing paralysis symptoms around 20-24 hours post injection. The experiment was terminated by the onset of symptoms, and after anesthetization using CO_2_ the queens’ abdomens were sampled and stored as described above.

For infection of the last group of reproductive queens, right after emergence, queens were caged and banked in a donor colony for one week. Then, they were taken out and treated with CO_2_ two times a week later and return to the donor colony for one more week to stimulate ovary development without mating (Cobey et al., 2013). Thereafter, queens were either injected with the IAPV-inoculum (n= 34), or PBS as sham control (n=31) and maintained with attendant workers in the incubator (35 °C, 65% humidity), as described above. Paralysis symptoms started in the IAPV-inoculated queens 40-44 hours post injection, when the experiment was terminated. Abdomens were sampled as described above.

### RNA isolation, cDNA Synthesis, and Quantitative PCR (qPCR) on experimentally infected queens

Each abdomen was transferred into a 2 mL bead ruptor homogenizing tubes containing 5 ceramic beads. The samples were then homogenized using an automated Bead Ruptor Elite (Omni International, Kennesaw, GA, USA) at speed of 5 meter per second (5 m/s) for 20 s. Followed by homogenization, total RNA was extracted using an established TRIzol™ (Invitrogen, Carlsbad, CA, USA) protocol. The concentration and purity of the extracted RNA samples were measured using a Nanodrop OneC Microvolume UV-Vis Spectrophotometer (Thermo Fisher Scientific, MA, USA). The total RNA concentration was adjusted to 20 ng/μL in molecular grade water (Fisher Scientific, Fair Lawn, NJ, USA). cDNA was synthesized using the High-Capacity cDNA Reverse-Transcription Kit (Applied Biosystems, Foster City, CA, USA). Ten microliters of the RNA template (200 ng) were added to 10 μL of the provided cDNA master mix, followed by an incubation period as recommended by the manufacturer; 10 min at 25 °C, 120 min at 37 °C, and 5 min at 85 °C. Resulting cDNA solution then diluted 10-fold in molecular grade water to serve as template in subsequent qPCR.

We quantified the IAPV intensity, expression level of three genes of interest including vitellogenin, protein lethal(2)essential for life-like, heat shock protein 70 cognate 4 (see ***Supplementary Table S7*** for primer sequences). Three housekeeping genes including actin, glyceraldehyde 3-phosphate dehydrogenase (GapDH) and ribosomal protein S5 (RPS5) were also amplified as internal control and for relative quantification.

The qPCR was conducted in duplicate using 384-well plates on a QuantStudio™ 6 cycler (Thermo Fisher Scientific). The reactions were performed using unlabeled primers and SYBR Green DNA binding dye (Applied Biosystems) with a volume of 12 μL and the final primer concentrations of 0.4 μM. We added RNase-free water as template for the NTC and no reverse transcriptase for the NRT controls. The thermal cycling conditions were 10 min at 95 °C, followed by 40 cycles consisting of a denaturing stage at 95 °C for 15 s and as annealing/extension stage at 60 °C for 1 min. This procedure was followed by a final melt-curve dissociation analysis to confirm the specificity of the products. For IAPV, the C_t_ values were determined at the same fluorescence threshold (0.05) for all plates, and a Ct value of 35 or lower was recorded as positive amplification. For three genes of interest and housekeeping genes we collected C_t_ values based on the default threshold as determined by the QuantStudio™ 6 for each target gene.

We quantified immune-gene expression for each sample using relative quantification. We calculated geometric mean of the C_t_ value from three reference genes (RPS5, actin, and GapDH) to confirm amplification. We subtracted the C_t_ value of geometric mean from target gene for each sample (ΔC_t_= C_t_(gene of interest) – C_t_(GMean of reference genes)). To calculate the ΔΔC_t_ for each target gene, we subtracted the average of the ΔC_t_ values across the control samples at each age from the ΔC_t_ for each target gene. Fold gene expression was then calculated using the 2^-(ΔΔCt)^ method. Pairwise comparisons of gene expression were evaluated using the wilcox.test function in the ggpubr package (v0.4.0). Data are plotted in the form of log_10_(fold gene expression) for clarity.

## Data Availability Statement

All data underlying the figures in this manuscript are supplied as supplementary information. The previously published (spermathecal fluid) raw data are publicly available on the MassIVE archive (www.massive.ucsd.edu, accession MSV000085428). The ovary mass spectrometry data are available on the MassIVE archive (www.massive.ucsd.edu, accession MSV000087150). Source code underlying figures and data analysis are available freely upon request.

**Supplementary Table S1** - complete sample metadata for the queens associated with Survey 3

**Supplementary Table S2** - ovary mass data for Surveys 1-3

**Supplementary Table S3 -** viral data for Survey 1

**Supplementary Table S4** - viral data for Survey 3

**Supplementary Table S5** - short term (2 d) queen stress ovary size data

**Supplementary Table S6** - long term (2 w) queen stress ovary size data

**Supplementary Table S7 -** primer sequence data used for viral and gene expression analysis

**Supplementary Table S8** - gene expression RT-qPCR data

**Supplementary File S1 -** Supplemental results & discussion, experimental heat-shock and pesticide stress applications, and confirmation of IAPV experimental infection using absolute qPCR.

## Conflicts of interest statement

The authors declare they have no conflicts of interest.

## Acknowledgements

Honey bee research in LJF’s group is supported by an NSERC Discovery Grant. Mass spectrometry infrastructure is supported by Genome Canada and Genome BC (264PRO) and computational infrastructure is supported by a Compute Canada Resource Allocation to LJF. AM was supported by an NSERC Postdoctoral Fellowship. A Project Apis m. grant to AM and LJF helped fund this research, as well as a Boone-Hodgeson-Wilkinson Trust Fund grant to AM and LJF. AC was supported by a Christi Heintz Memorial Award. A part of this research was performed while EA held an NRC Research Associateship award and supported by the US Army Research Office (grant # W911NF1520045) and US Department of Agriculture-APHIS (AP20PPQS&T00C014). We would like to acknowledge Hives for Humanity, Robyn Underwood, and numerous beekeepers in the BC Bee Breeders’ Association for their contribution to our queen surveys.

## Author contributions

AC and AM wrote the first draft of the manuscript, made the figures, and interpreted the data with editing assistance from LJF and DRT. AC and AM conducted queen survey 1, and AM conducted queen surveys 2 and 3 with assistance from acknowledged collaborators. AC conducted ovary proteomics and data analysis. AM conducted the experiments involving alternate abiotic stressors. AC and AM conducted the IAPV infection experiment for ovary mass. EM conducted the qPCR analysis of viruses in queens from survey 3. Grants supplied to AC, AM, LJF, and DRT funded the queen survey and proteomics work. Grants to EA, OR, and BH funded the experimental infection and associated qPCR analysis. EA and BH conducted the infections, associated qPCRs and data analysis. OR, EA, and BH conceptualized the experimental queen infections. AM and AC conceptualized the queen survey study and associated proteomics analyses.

## Notes

### Competing Interest Statement

The authors have declared no competing interest.

### Summary of Updates

Added new data, updated figures, added new references, and reduced focus on the reproduction-immunity trade-off hypothesis

