## Supplementary File S1 for "Fertility costs of cryptic viral infections in a model social insect"

**Supplementary Results & Discussion**

*Short-term temperature and pesticide stress do not reduce ovary mass*

Heat stress induces expression of heat-shock proteins in queens^1,2^, is known to damage stored sperm^2-4^, and has been linked to queen failure^5,6^. Most research has focused on the damaging effects of heat on sperm viability, but theoretically the energetic expense of upregulating heat-shock protein expression could be sufficient to reduce the mass of ovaries, even in the absence of infection. Therefore, it is possible that the link between ovary size and queen failure could be driven by temperature stress alone, with viral infection playing a lesser role. To test this hypothesis, we retrieved n = 19 queens from their colonies, heat-shocked n = 10 queens (2 h at 42°C), and incubated the remaining n = 9 queens at hive temperature (33°C, 2 h) prior to returning them to their respective colonies. We measured their ovary masses two weeks after the heat-stress but found no significant differences (linear model; F = 1.12; df = 1, 17; p = 0.31; **Figure S1 a-b, Supplementary Table S3**).

It is possible that two weeks is sufficient time for a drop in ovary mass to subsequently rebound; therefore, we also conducted a laboratory experiment in which queens were exposed to heat (42°C) for 0 or 2 h and allowed to recover for two days at 33°C prior to ovary dissection, using queens from two different sources (**Figure S1 b**; California queens for Experiment 1, and locally raised queens for Experiment 2). We found no significant effect of heat shock across experiments (F = 0.055, df = 1, 44, p = 0.82). Together, these data show that the heat-shock response alone is not sufficient to deplete the resources available to invest in ovary size, and suggests that the negative relationship between viral infection and ovary mass is more likely driven by a specific effect of the viruses. Data associated with these experiments are available in **Supplementary Table S4**.

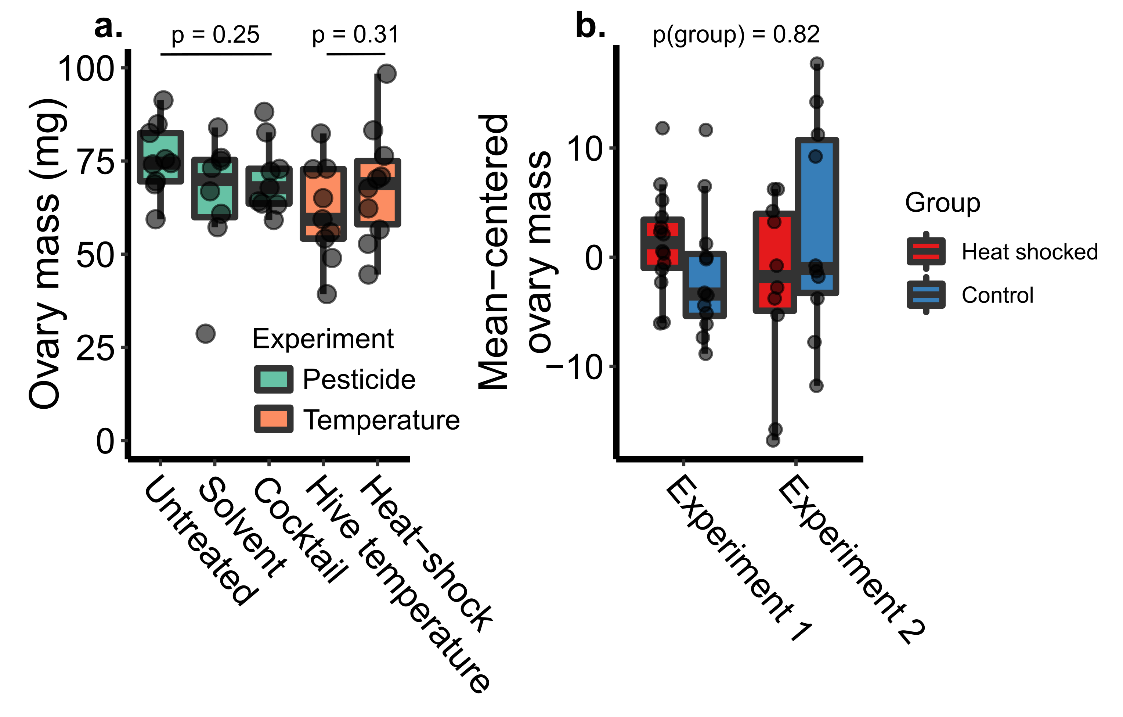

**Figure S1.** *Impact of heat stress and pesticide stress on queen mass and ovary mass.* For the pesticide stress experiment, we randomly assigned queens to either untreated (n = 9), solvent-treated (n = 8), or pesticide-treated (n = 9) groups. In a second experiment, we randomly assigned queens to either heat-shocked (42°C, n = 10) or hive temperature control (33°C, n = 9) groups. Boxes represent the interquartile range, bars indicate the median, and whiskers span 1.5 times the interquartile range.

Pesticide stress is an additional environmental stressor faced by all bees including queens, and it is a second potentially confounding, unknown variable that may have influenced ovary mass in our queen survey. We previously found that failed queens express higher levels of proteins associated with detoxification^1^. Furthermore, high levels of pesticides within wax have been linked to ‘queen events’ (supersedure or queen loss) in US beekeeping operations^7^. It is possible that the energetic cost of combating pesticide stress could also lead to a reduction in ovary mass and subsequent poor performance. With a different set of n = 26 queens, we repeated the same procedure for pesticide stress as for heat stress. We administered queens with a topical dose of mixed agrochemicals frequently found in wax, as reported by Traynor *et al.*^7^, and which we have previously utilized^1^ (diluted in acetone; n = 9), an acetone control (n = 8), or no treatment (n = 9). We used a dose equivalent to a hazard quotient (cumulative toxicity of each mixture component) of 3,500, which is similar to the level detected in wax of colonies experiencing queen events, and is ~7-fold higher than the level at which we have previously detected putative detoxification responses by proteomics^1^. We again found no relationship between ovary mass and treatment group (linear model; F=1.49; df = 2, 23; p = 0.24), suggesting that the stress associated with this pesticide cocktail is not sufficient to divert resources away from ovary investment. Data associated with these experiments are available in **Supplementary Table S4**.

While pesticide stress has previously been associated with queen events^7,8^, we did not observe a consistent pattern of queen events associated with treatment group: two attempted supersedures occurred in the untreated control, one attempted supersedure and one queen was lost in the acetone group as well as the pesticide group. It is possible that we did not detect a relationship between queen events and groups may be because this was not a long-term, chronic pesticide stress as queens experience in the field. It is also possible that queen events occur due to indirect effects of pesticide stress on workers^9,10^, which would not be detectible by our experimental design since the queens were exposed directly.

*Confirmation of IAPV infection in experimental innoculations*

We infected queen pupae, two day old adult queens and two week old adult queens via injection as described in the Methods, and confirmed the presence of infection using RT-qPCR (**Figure S2**).

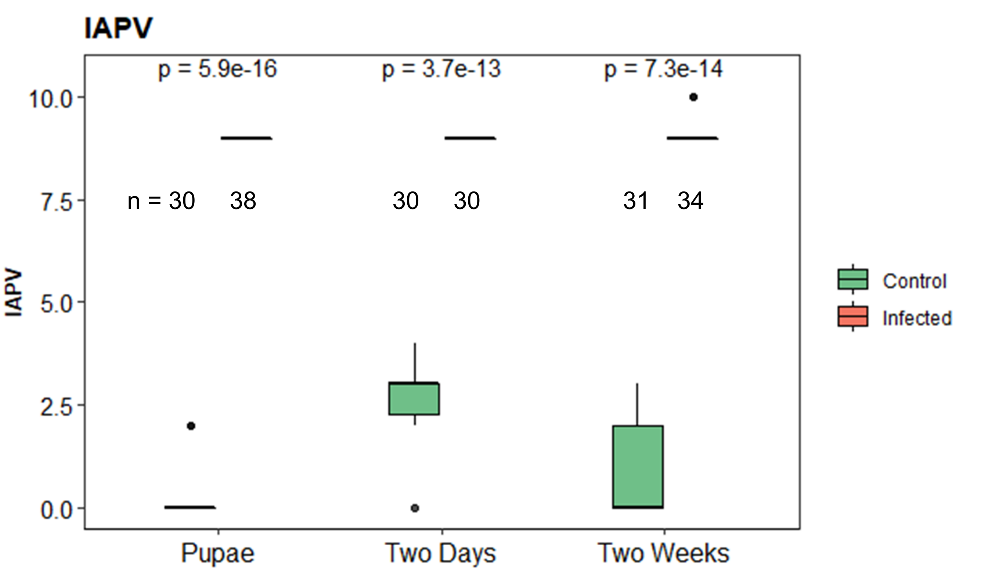

**Figure S2.** *Confirmation of IAPV infection.* IAPV load in each sample was quantified using absolute quantification, based on our standard curves obtained through serial dilutions of known numbers of amplicons as described before^11^. To improve data compliance with parametric assumptions, statistical tests were performed on transformed raw data, according to *x*′=*log*10(*x*+1). The y axis indicates log10 transformed viral RNA copies.

*References*

1	McAfee, A. *et al.* Candidate stress biomarkers for queen failure diagnostics. *BMC Genomics* **21**, 571, doi:10.1186/s12864-020-06992-2 (2020).

2	McAfee, A. *et al.* Vulnerability of honey bee queens to heat-induced loss of fertility. *Nature Sustainability*, 1-10 (2020).

3	Pettis, J. S., Rice, N., Joselow, K., vanEngelsdorp, D. & Chaimanee, V. Colony Failure Linked to Low Sperm Viability in Honey Bee (Apis mellifera) Queens and an Exploration of Potential Causative Factors. *PLoS One* **11**, e0147220, doi:10.1371/journal.pone.0147220 (2016).

4	Rousseau, A., Houle, É. & Giovenazzo, P. Effect of shipping boxes, attendant bees, and temperature on honey bee queen sperm quality (Apis mellifera). *Apidologie*, 1-12 (2020).

5	Withrow, J. M., Pettis, J. S. & Tarpy, D. R. Effects of temperature during package transportation on queen establishment and survival in honey bees (Hymenoptera: Apidae). *Journal of economic entomology* **112**, 1043-1049 (2019).

6	Guarna, M. M., Pettis, J. S. & Pernal, S. F. in *EurBee 8, 8th Congress of Apidology.* 132.

7	Traynor, K. S. *et al.* In-hive Pesticide Exposome: Assessing risks to migratory honey bees from in-hive pesticide contamination in the Eastern United States. *Scientific Reports* **6**, 33207 (2016).

8	Tsvetkov, N. *et al.* Chronic exposure to neonicotinoids reduces honey bee health near corn crops. *Science* **356**, 1395-1397, doi:10.1126/science.aam7470 (2017).

9	Milone, J. P. & Tarpy, D. R. Effects of developmental exposure to pesticides in wax and pollen on honey bee (Apis mellifera) queen reproductive phenotypes. *Sci Rep* **11**, 1020, doi:10.1038/s41598-020-80446-3 (2021).

10	Milone, J. P., Chakrabarti, P., Sagili, R. R. & Tarpy, D. R. Colony-level pesticide exposure affects honey bee (Apis mellifera L.) royal jelly production and nutritional composition. *Chemosphere* **263**, 128183 (2020).

11	Francis, R. M., Nielsen, S. L. & Kryger, P. Varroa-virus interaction in collapsing honey bee colonies. *PLoS One* **8**, e57540 (2013).
